# Systems genetics identifies the association between Enc1 and cognitive function in the hippocampus

**DOI:** 10.1101/2023.10.23.563569

**Authors:** Hongjie He, Ran Tao, Zhe Han, Quanting Yin, Shuijing Pan, Lu Lu, Akhilesh Kumar Bajpai, Jia Mi, Donglai Qi, He Li, Fuyi Xu

## Abstract

Ectodermal-Neural Cortex 1 (ENC1) is expressed in multiple regions of the brain, including the hippocampus. However, knowledge about its function has been well explored only in the context of peroxidative stress and cancer. In this study, we investigated the association of hippocampal Enc1 with cognitive function in BXD mice. We performed Pearson correlation, phenotype-wide association analysis (PheWAS), expression-Based PheWAS, pathway enrichment, and protein interaction networks on Enc1 and BXD phenotypes/transcriptome of the hippocampus, and the results indicated that Enc1 is inextricably linked to cognitive performance. In addition, we found that most of the *Enc1* co-expressed genes were highly expressed in GABAergic neuronal cells. Expression quantitative trait loci analysis indicated that Enc1 was *cis*-regulated in the hippocampus of mice as well as human. Genome-wide association analysis revealed ENC1 to be significantly associated with cognitive-related traits, including age-related cognitive changes etc. In conclusion, our findings demonstrated that *Enc1* is involved in cognitive functions mainly in hippocampal GABAergic neuronal cells through neurogenesis, synaptic signaling, and CGMP-PKG signaling pathways, and interacts with the neurological function-related genes.

## Introduction

Cognition, the process of acquiring or applying knowledge, or the process of information processing, is the most basic human mental process, including sensation, perception, memory, thinking, imagination, and language. Cognitive dysfunction refers to various degrees of cognitive damage due to various causes, ranging from mild cognitive impairment to dementia[1]. In 2019, an estimated 55.2 million people worldwide suffer from dementia and 1.6 million die from it, making it to be the seventh leading cause of death, which brings a serious global health burden[2]. Currently, dozens of genes associated with cognition have been identified, including APOE[3], MEF2[4], COMT[5], KMO[6], BRCA2[7], etc., leading to advances in the treatment of cognitive disorders. However, due to the complexity of cognitive dysfunction, the role of common genetic variants and the pathogenic mechanisms have not been fully elucidated. Therefore, the identification of new causative genes and drug targets may provide new ideas and approaches for the diagnosis, treatment and prevention of cognitive impairment.

*Enc1* (Ectodermal-Neural Cortex 1) encodes an actin-binding protein, a member of the KELCH family. It has primarily been investigated for its role as a regulator of the transcription factor Nrf2 in the oxidative stress response[8]. Previous studies have also shown that it plays a key role in neuronal and adipocyte differentiation, and is involved in many biological processes in cancer, including ovarian cancer[9], endometrial cancer[10], and lung cancer[11]. Interestingly, it is highly expressed in brain tissues including the frontal cortex, cortex and anterior cingulate cortex, and hippocampus compared to other tissues (GTEx, https://gtexportal.org/home/). In recent years, ENC1 has been found to be associated with residual cognition by genome-wide association analysis[12]. Differential expression analysis revealed that it is down-regulated in the brains of Alzheimer’s disease (AD) patients, and It has been further thought to be associated with the development of AD through gene function enrichment analysis and protein-protein interaction (PPI) network construction[13,14]. ENC1 knockdown in SH-SY5Y cells (human neuroblastoma cells) alleviates death of Huntington protein-expressing neuronal cells under ER stress[15]. However, it remains unclear about whether Enc1 is involved in cognitive regulation and what the biological mechanisms are.

Systems genetics is an approach to understanding the flow of biological information underlying complex traits. It considers not only underlying genetic variation but also intermediate phenotypes, including gene, protein and metabolite levels, as well as gene-by-gene and gene-by-environment interactions[16,17]. Due to the complexity, environmental uncontrollability, and ethical issues, cognitive-related disorders are difficult to study directly in human populations. Studies of genetic reference populations (GRPs) in mice with controllable environmental factors provide a stable platform for identifying genetic, environmental, and GxE factors that influence such complex traits. Among the current animal GRPs, the BXD mouse panel, generated from the progeny of crosses between C57BL/6J (B6) and DBA/2J (D2) strains after more than 20 consecutive generations of sibling self-fertilization, serves as a suitable animal model[18,19]. This family contains over 200 strains with ∼150 whole-genome sequenced, and segregates for 6 million common DNA variants[18]. In addition, the B6 and D2 parental strains showed significant differences in model separation of cognitive abilities[20], reversal learning[21], and several types of complex learning including contextual fear conditioning[22]. Furthermore, genetic differences between them have been shown to translate into a wide range of CNS-related functional and molecular associations, including hippocampus-related memory and learning tasks, and postsynaptic and presynaptic protein expression[23].

In this study, we collected phenotypic and hippocampal transcriptomic data of BXD mice, as well as publicly available genomic data of humans, and investigated the effects of hippocampal *Enc1* on cognition through a combined approach of systems genetics and bioinformatics analyses.

## Materials and methods

### Hippocampus transcriptomic dataset

The hippocampus is a vital part of the mammalian central nervous system that plays a role in short-term and long-term memory, as well as spatial orientation, and is associated with various cognitive deficit disorders. The Hippocampus Consortium data set in this study provides estimates of mRNA expression in the adult hippocampus of 71 genetically diverse strains of mice, including 67 BXD recombinant inbred strains, their parental strains (C57BL/6J, DBA2/J), and two reciprocal F1 hybrids. The standardized dataset “hippocampus consortium” is available on GeneNetwork (https://gn1.genenetwork.org/) and belongs to the group “BXD Family” and the type “hippocampus mRNA” with the dataset name “hippocampus Consortium M430v2 (June 6) RMA” (GEO Series: GSE84767). The following is a brief description of the process used for generating this dataset.

### Mice and tissue harvesting

BXD mice were obtained from the University of Tennessee Health Science Center (UTHSC), University of Alabama (UAB), or directly from the Jackson Laboratory and were housed in UTHSC, Beth Israel Deaconess, or the Jackson Laboratory prior to execution. The BXD mice in this study were from 71 strains, and the vast majority were between 45 and 90 days old (mean 66 days, maximum range 41 to 196 days). All the animals were executed by cervical dislocation between 9 a.m. and 5 p.m. The brain was then removed, placed in an RNAlater solution, and the entire hippocampus dissected. The cerebellum and olfactory bulb were removed; the brain was semi-excised and both hippocampal bodies were dissected as a whole.

### RNA extraction and evaluation

Anatomical tissues from six hippocampi and three naive adults of the same strain, sex, and age were collected in one session and used to generate cRNA samples. The 206 RNA samples were extracted with RNA STAT-60 according to the manufacturer’s instructions, which mainly included tissue homogenization, RNA extraction, RNA pretreatment, RNA washing, and RNA purification. Finally, RNA purity was evaluated using the 260/280 nm absorbance ratio, and values had to be greater than 1.8. The Agilent Bioanalyzer 2100 was used for assessing RNA integrity and samples with an RNA integrity number (RIN) greater than 8 were considered.

### Microarray and data normalization

Mixed RNA samples from 2 to 3 animals from the same age, strain and sex were hybridized to a single Affymetrix GeneChip Mouse Expression 430 2.0 short oligomer array. The Affymetrix 430v2 arrays consist of 992936 useful 25-nucleotide probes that can estimate the expression of approximately 39,000 transcripts and most known genes and expressed sequence tags. The raw microarray data were normalized using the robust multi-array average (RMA) method[24] and further converted to an improved z score (2z+8). This analysis was performed using the R statistical function. Additional details on RNA extraction and hybridization protocols can be found in NCBI-GEO under the accession GSE84767.

### Correlation analysis

Pearson correlation analysis was used to identify *Enc1* co-expressed genes or published traits/phenotypes associated with the gene being explored. P-values less than 0.05 were considered statistically significant. This analysis was performed on GeneNetwork[25].

### Phenome-wide association analysis (PheWAS)

PheWAS is a reverse genetic analysis method to identify the potential phenotypes associated with genetic variations[26]. Genes that contain high-impact variants, including missense, nonsense, splice site, frameshift mutations, copy number variations (CNVs), as well as genes that have significant cis-e(p)QTLs in the BXD transcriptome and proteome datasets were included in the PheWAS analysis. Genetic variants of each gene are represented by the SNPs within the genes as well as their cis-QTLs. About 5,000 clinical phenotypes were used to study the association between genes and phenotypes. In this paper we used a multi-locus mixed-model approach (mlmm) to estimate the associations between each gene (represented by the genetic variants of the gene) and clinical traits. This analysis was performed on Systems Genetics at EPFL (https://systems-genetics.org)[27].

### Expression-Based PheWAS (ePhewas)

Associations between transcripts/proteins and phenotypic traits were estimated using mixed model regression analysis. Transcript-trait pairs with fewer than 15 overlapping lines were removed from the analysis. Phenotype-wide significance analysis was performed using Bonferroni correction. This analysis was performed on Systems Genetics at EPFL (https://systems-genetics.org)[27].

### Expression quantitative trait loci (eQTLs)

In the analyses presented here, genome-wide eQTL mapping was performed using 71 BXD strains on the GeneNetwork using a modified Haley-Knott regression method as well as the GEMMA method. GEMMA corrects for kinship between samples using a linear mixed model approach and allows the user to fit multiple covariates such as sex, age, treatment, and genetic markers[28]. It incorporates the Leave One Chromosome Out (LOCO) method to ensure that correction for relatedness does not eliminate useful genetic variation in the vicinity of each marker, and association results are presented in –log10P. The likelihood ratio statistic (LRS) generated by the Haley-Knott regression method is used to assess associations between genotype and gene expression levels[29]. The threshold for genome-wide significance was determined by 1,000 permutation tests. This analysis was performed on GeneNetwork.

### Sequence variants

Genetic variations between parental strains B6 (C57BL/6J) and D2 (DBA/2J) were searched with MGI (Mouse Genome Informatics, https://www.informatics.jax.org/).

### PPI Network

PPI networks were constructed and evaluated with NetworkAnalyst (https://www.networkanalyst.ca/)[30], which uses comprehensive data collated from the International Molecular Exchange (IMEx) compliant literature curated database, InnateDB[31]. InnateDB (http://www.innatedb.com) is an integrated analysis platform dedicated to facilitating systems-level analysis of mammalian innate immune networks, pathways, and genes in IMEx, which is dedicated to developing rules for describing molecular interaction data and actively curating these interactions from the scientific literature[31].

### Pathway enrichment analysis of co-expressed genes

WEB-based Gene SeT AnaLysis Toolkit (WebGestalt) (http://bioinfo.vanderbilt.edu/webgestalt/) was used for Gene Ontology Biological Processes (GO-BP), the Kyoto Encyclopedia of Genes and Genomes (KEGG) Pathways and Mammalian Phenotype Ontology (MPO) enrichment analysis[32]. FDR lower than 0.05 is considered statistically significant.

### Exploration of Gene Function

To identify genes with functions associated with cognition, we used “cognition” as a keyword and retrieved genes from the Mouse Genome Informatics (MGI)[33] phenotype/disease association (Human-Mouse: Disease Connection) (https://www.informatics.jax.org/), the NHGRI-EBI GWAS Catalog[34] (https://www.ebi.ac.uk/gwas/), and International Mouse Phenotyping Consortium (IMPC)[35] (https://www.mousephenotype.org/). Translation of human gene lists from MGI and GWAS to mouse homologs was done using the homologene R package (https://github.com/oganm/homologene).

### Cell type composition of *cis*-eQTL genes in the brain

We used Mouse Cell Atlas (MCA) (https://bis.zju.edu.cn/MCA/) database to retrieve the tissue type “Adult Brain” and obtained raw data on the expression of each gene in the barcodes of 15 brain cell types[36], and by “averaging”, we investigated in which cell types the *Enc1* co-expressed genes were highly expressed. Normalization was performed using Z-score method in the pheatmap R package, using clustering by row.

## Results

### Enc1 mRNA levels in the hippocampus significantly correlate with cognition-related properties

To investigate the effect of *Enc1* on cognition, we first determined its expression (1450061_at, a probe located in the distal 3’UTR) in the hippocampus across 71 BXD strains. The average *Enc1* expression in the hippocampus was 13.51 ± 0.11 SD, with the highest and lowest expressions being 13.77 in BXD86 and 13.16 in BXD87 (**Fig. A.1**). Next, we conducted correlation analysis and found *Enc1* to be significantly associated (p < 0.05) with 150 nervous system-related traits. For instance, *Enc1* was positively correlated with the rate of unsuccessful spontaneous alternations in the y-maze, negatively correlated with the pre-stimulus freeze time in the situational fear experiment, the number of touches on the lever in the sucrose self-administration experiment, and the number of touch screens in the object localization paired-association learning touch screen experiment (**Fig. B.1, Table 1**). Furthermore, PheWAS between *Enc1* genotypes and BXD phenotypes (including about 5000 traits) revealed 35 nervous system-related traits exhibiting moderate associations, including neuron number, anxiety-fear response, spatial navigation (-log10(p) >= 2, **Fig. C.1**). The ePheWAS analysis of *Enc1* gene mRNA expression in the hippocampus with BXD phenotypes also yielded 13 moderately associated neurological related phenotypes (-log10(p) >= 2, **Fig. D.1**), such as the object localization paired-association learning, dopamine receptor binding. Further, *Enc1* knockdown causes changes in cholesterol, HDL cholesterol, bilirubin, alkaline phosphatase levels and glucose levels from IMPC database, which have been associated with cognitive dysfunction (**Table 2**).

**Table 1.**
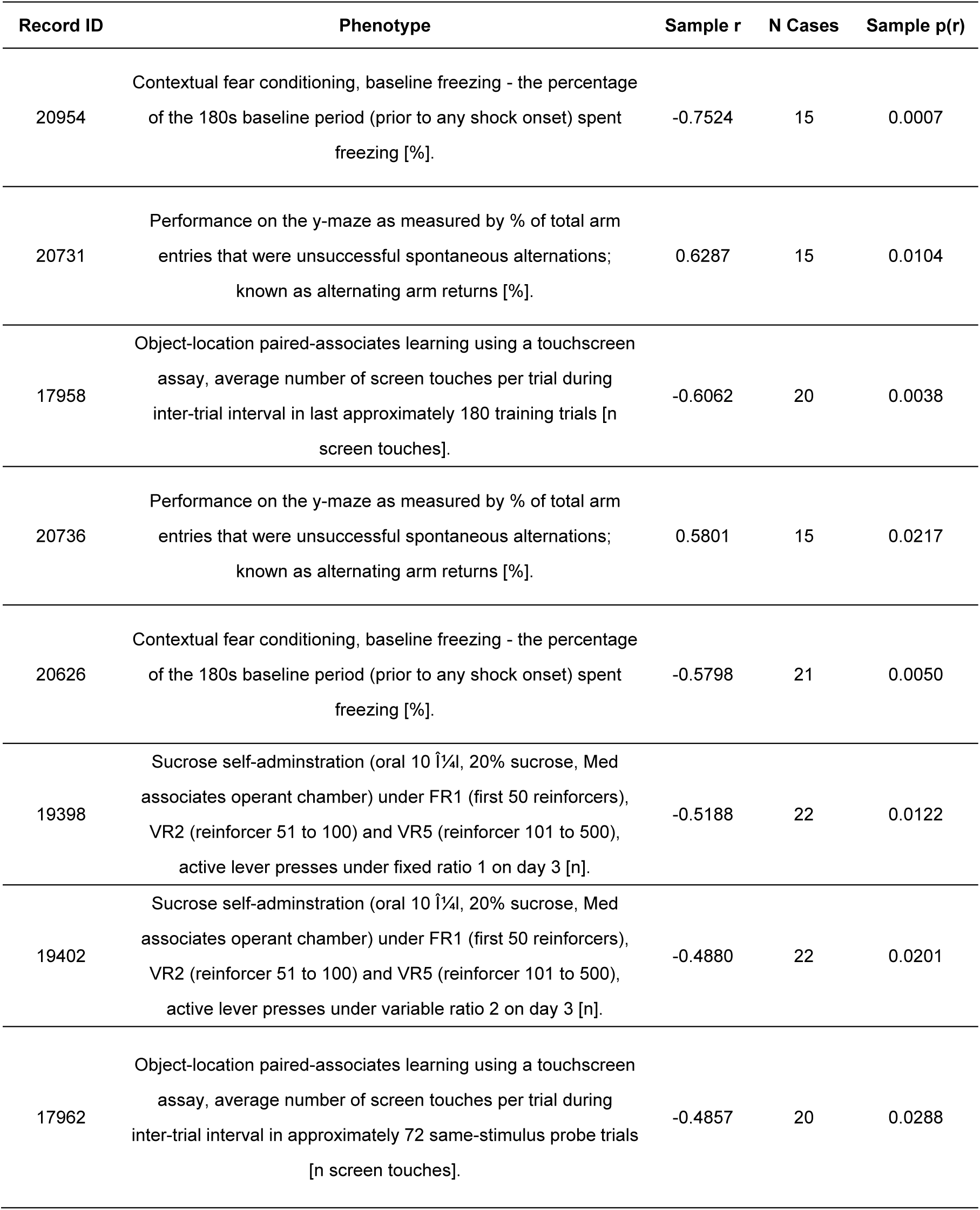
Correlation of *Enc1* with cognition relevant phenotypes. Notes: More details for the phenotype descriptions can be access on genenetwork via record ID.

**Table 2.**
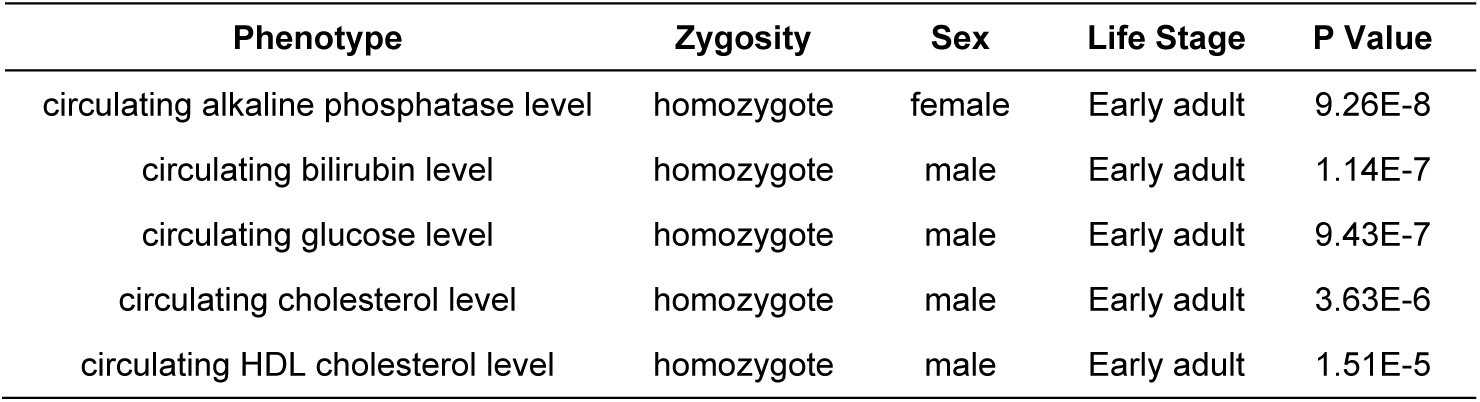
Lists of altered metabolic traits in Enc1 knockout mice. Notes: Data was obtained from the IMPC (International Mouse Phenotyping Consortium, https://www.mousephenotype.org/).

### Genetic correlations between *Enc1* and hippocampal transcriptome

To gain insight into the pathways and biological functions of *Enc1* involved in the hippocampus, we performed a Pearson genetic correlation analysis of *Enc1* with the entire hippocampal transcriptome. Intersection statistics were then performed between the top 2000 *Enc1* correlated probes (corresponding to 1841 transcripts, p<0.05) and the cognition related genes generated from the IMPC (485 genes), GWAS (1598 genes), and MGI (1467 genes). Results revealed a total of 308 *Enc1* correlated genes were implicated in cognition related functions (**Fig. A.2**). Three genes, *Cnksr2*, *Mbd5*, and *Gria1*, were shared across four gene sets and were positively correlated with *Enc1* (**Fig. B-D.2**). *Cnksr2* (connector enhancer of kinase suppressor of Ras 2) is expressed in cortical, striatal and cerebellar regions and is localized at excitatory and inhibitory post-synapses. *Cnksr2* knockout mice exhibit marked anxiety, impaired learning and memory, and progressive and dramatic loss of ultrasonic vocalizations[37]. *Mbd5* (methyl-CpG-binding domain protein 5) mouse carrying an insertional mutation in *Mbd5* exhibits abnormal social behavior, cognitive impairment, and motor and craniofacial abnormalities[38]. *Gria1* (Glutamate Ionotropic Receptor AMPA Type Subunit 1) encodes the GluA1 subunit of α-amino-3-hydroxy-5-methyl-4-isoxazole propionate (AMPA) receptors, which are ligand-gated ion channels that act as excitatory receptors for the neurotransmitter L-glutamate (Glu). The Xenopus *Gria1* models show transient motor deficits, an intermittent seizure phenotype, and a significant impairment to working memory in mutants[39].

### PPI Network Analysis infers key regulators for *Enc1* co-expressed genes in the hippocampus

To investigate the key regulators in *Enc1* co-expressed genes, we submitted 1841 transcripts and *Enc1* to NetworkAnalyst and evaluated the PPI network with curated and non-redundant sets of protein interactions from the IMEx consortium database. The PPI network showed that 20 genes (*Ubc, Ppp2ca, Mapk1, Ikbkb, Myod1, Foxp3, Sall4, Nphp1, Irf4, Myd88, Rnf2, Ndn, Nfkb1, Plcg2, Traf6, Crep1, Ppp1cc, Bcl2, Ywhae, Pmp*) are at the central nodes of the network and have the most connections with other genes. Among them, *Ubc* (Ubiquitin C) had the highest connectivity with other genes and was located in the central hub position, followed by *Ywhae*, *Rnf2*, *Traf6* (**Fig. 3**). Notably, most of nodal genes are involved in the regulation of cognitive and neurological related traits. For instance, *Ubc* gene is a stress-protective gene[40]. Increased oxidative stress is inextricably linked to cognition[41]. In addition, *Ubc* is mediated by HSF1 (heat shock factor 1), which acts as a transcription factor and plays a central role in protein homeostasis, establishment and maintenance of synaptic fidelity and function, and memory consolidation[42].

**Figure 1.**
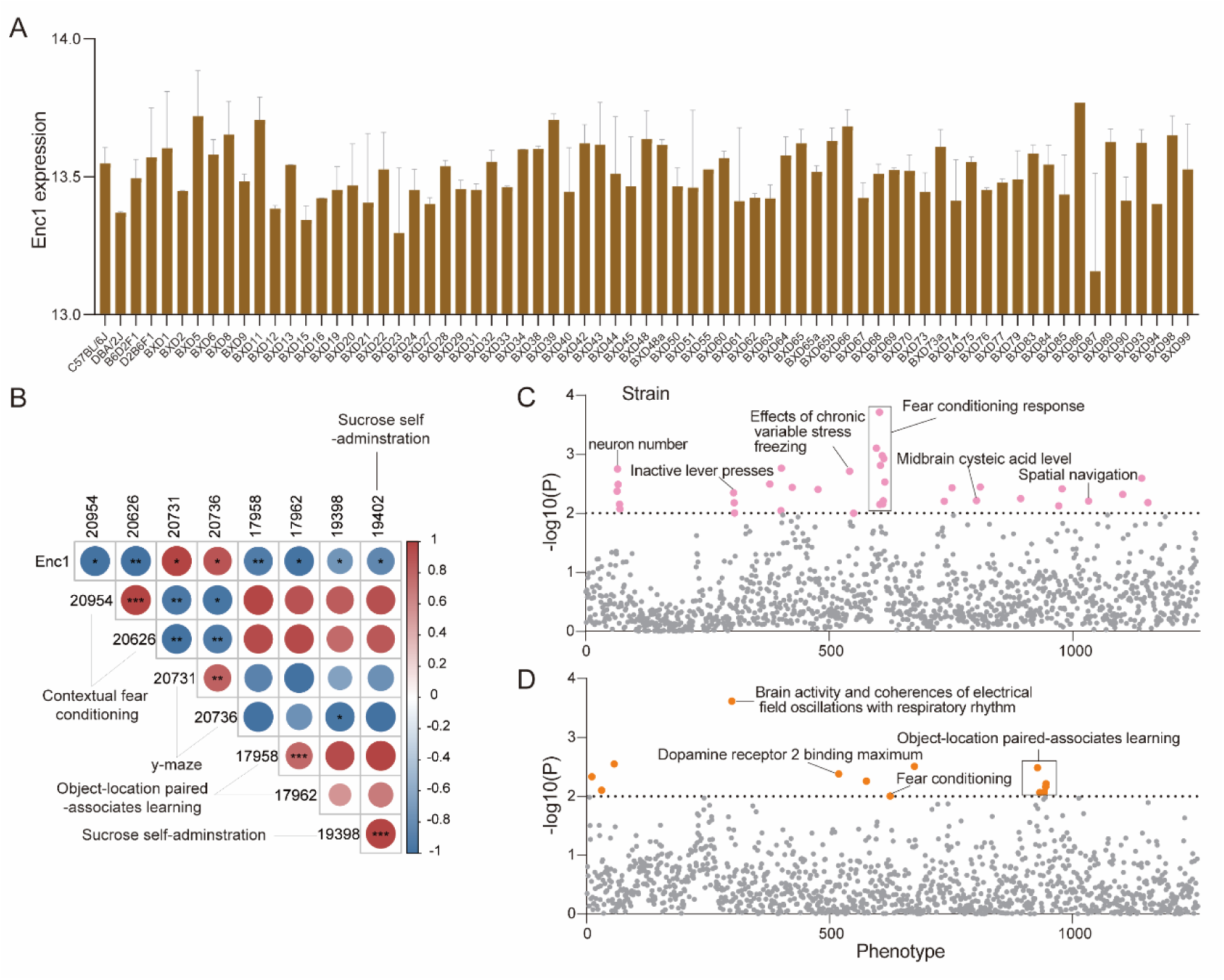
– *Enc1* expression in the hippocampus was associated with cognitive phenotypes in the BXD mice. (A) mRNA expression (mean ± SE) of *Enc1* in the hippocampus of 69 BXD strains and two parental strains (B6 and D2). The values are log2 transformed. (B) Correlation heatmap between *Enc1* and five cognition related traits (y-maze, contextual fear conditioning, Object-location paired-associates learning, Sucrose self-administration). The color and size of the circle represents the Pearson correlation R value, with red and blue representing positive and negative correlations, respectively. *p < 0.05, **p < 0.01, ***p <0.001. (C-D) Manhattan plots of PheWAS and ePheWAS showing the *Enc1*-associated phenotypes in BXD mice. Each point represents a neurological phenotype with a threshold line of –log10p = 2.

**Figure 2.**
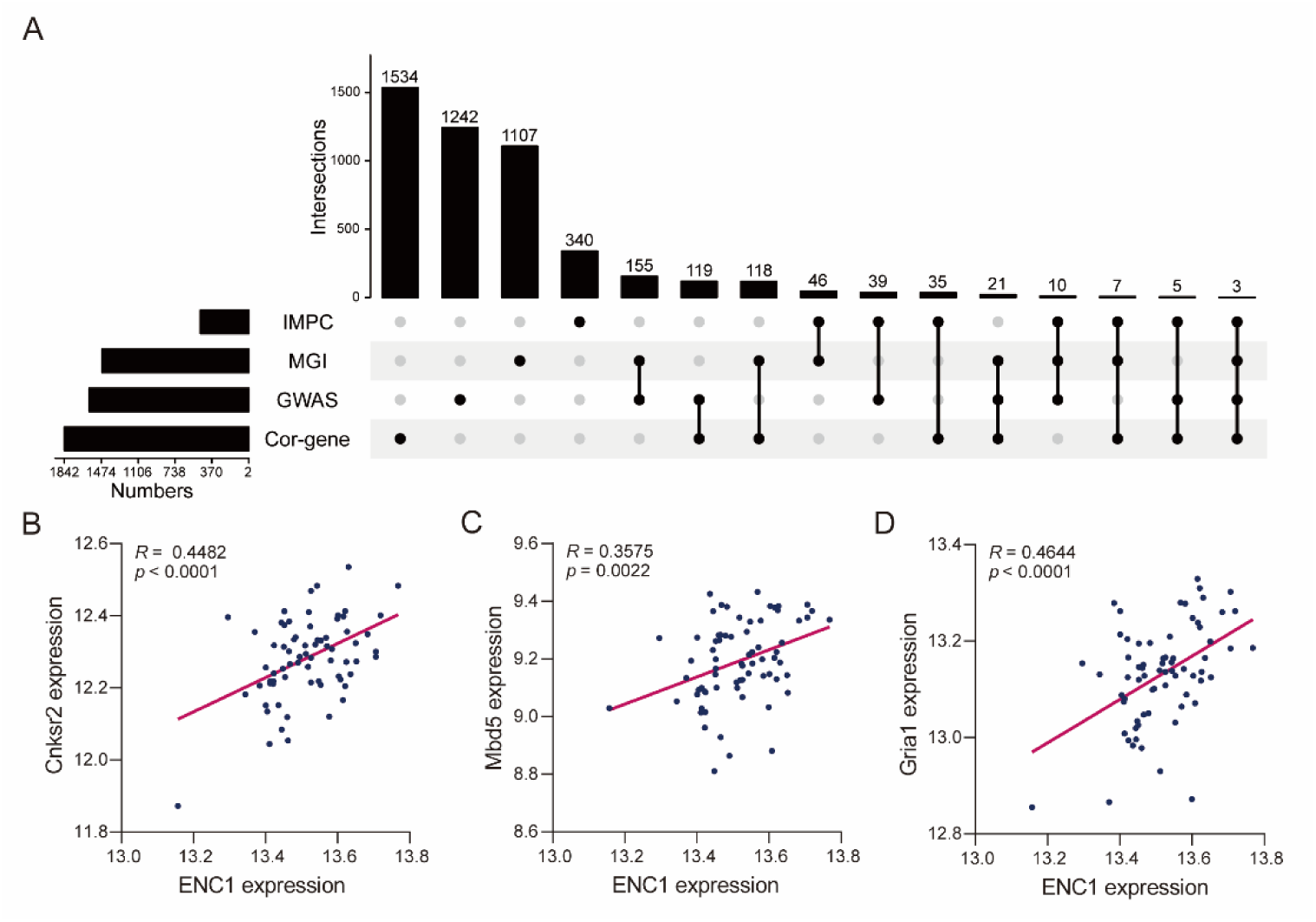
– Exploration of cognition related genes in *Enc1* co-expressed genes. (A) Intersections between *Enc1* co-expressed genes and cognition related genes generated from MGI, IMPC, and GWAS. The bars on the left represent the number of genes in each gene set, and the dotted line shows the gene sets that correspond to the number of overlapping genes in the bars above. MGI, Mouse Genome Informatics; IMPC, International Mouse Phenotyping Consortium; GWAS, GWAS Catalog; Cor_gene, *Enc1* correlated genes. (B-D) Correlations between *Cnksr2*, *Mbd5*, *Gria1* genes and *Enc1* gene expression. Each point represents a BXD strain. Pearson correlation coefficient is in the upper left corner and is significant at p<0.05.

**Figure 3.**
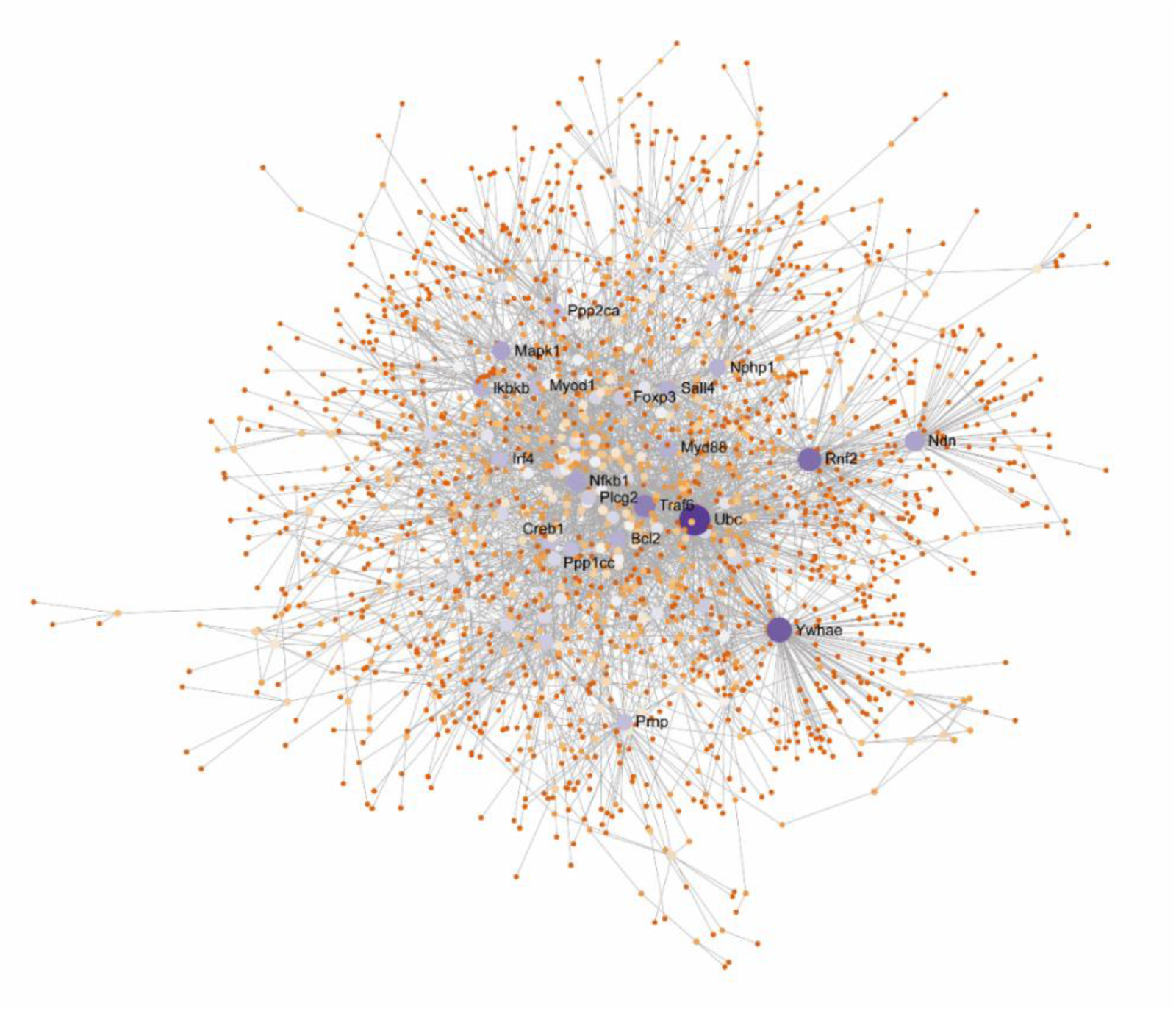
– PPI network analysis inferred key regulators of *Enc1* co-expressed genes. The Generic PPI network was constructed using the IMEx Interactome on NetworkAnalyst (https://www.networkanalyst.ca/) with *Enc1* co-expressed genes as input. The nodes in the network represent genes. Key node genes are indicated with gene symbols.

### *Enc1* co-expressed genes are involved in cognitive function

To investigate the specific biological processes involved in cognition, we used *Enc1* and its co-expressed genes for performing GO, KEGG and MPO enrichment analysis. GO enrichment showed that co-expressed genes are significantly enriched in neurogenesis and signaling related terms (**Fig. A.4**), such as neurogenesis (GO:0022008), synaptic signaling (GO:0099536) and generation of neurons (GO:0048699). In the neurogenesis process, 182 genes were significantly enriched, and the PPI network of these genes showed *Ywhae* as the hub gene, followed by *Ndn, Mapk1, Nphp1, Bcl2, Pax3, Rac1, App, Ppp1cc* that acted as the nodal genes (**Fig. B.4**). Microdeletion of chromosome 17p13.3 harboring *YWHAE,* presents with growth restriction, craniofacial dysmorphism, structural brain abnormalities and cognitive impairment[43]. KEGG enrichment showed that *Enc1* co-expressed genes are not only associated with signaling and synaptic transmission (**Fig. C.4**), such as cGMP-PKG signaling pathway (mmu04022), Dopaminergic synapse (mmu04728), but also significantly involved in cognitive and memory related pathways (**Fig. C.4**), such as AD (mmu05010), and Long-term potentiation (mmu04720). Thirty-two genes were enriched in the AD pathway, with *Mapk1*, *App*, *Calm1*, *Snca*, *Itpr1*, and *Bad* genes as the nodal genes (**Fig. D.4**). Inhibition of MAPK1 expression improves cognitive function in AD rats[44]. In addition, the enrichment of mammalian phenotype ontology (MPO) suggested that these genes are not only associated with learning, cognition, memory, and movement aspects (**Fig. E.4**), such as abnormal spatial learning (MP:0001463), abnormal learning/memory/conditioning (MP:0002063), abnormal cognition (MP:0014114), but are also significantly involved in synaptic transmission-related pathways (**Fig. E.4**), such as abnormal synaptic transmission (MP:0003635), abnormal CNS synaptic transmission (MP:0002206). A total of 97 genes were significantly enriched in the abnormal synaptic transmission pathway, with *App*, *Prnp*, *Nodd4*, *Creb1*, and *Snap25* as the critical node genes (**Fig. F.4**). Mutations in the *App* (amyloid-β precursor protein) gene, a key gene in all 3 of these pathways, involves or influences the process of amyloid Aβ production. Age spots formed by amyloid β protein are a characteristic lesion of AD. According to the amyloid cascade hypothesis, Aβ oligomers trigger vascular lesions, synaptopathy and glial activation[45]. In addition, a study has demonstrated that a coding mutation (A673T) in the APP gene prevents AD and cognitive decline in older adults without AD[46].

**Figure 4.**
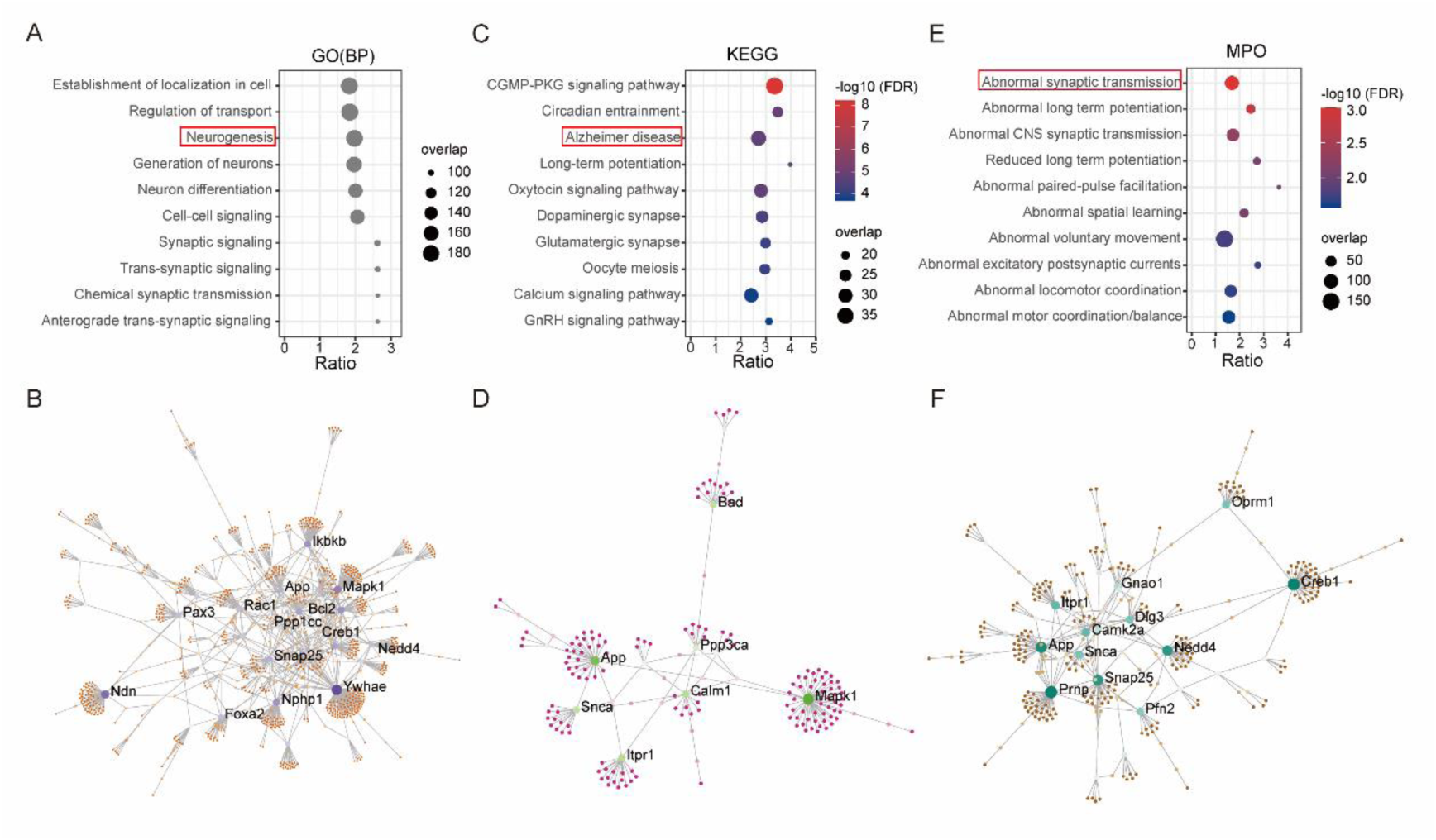
– Enrichment of processes, pathways, and mammalian phenotype ontologies by *Enc1* and its co-expressed genes. Bubble diagrams of (A) GO, (C) KEGG pathway and (E) MPO. Top ten terms of each enrichment are shown (FDR<0.05). The X-axis represents the enrichment ratio, whereas the Y-axis represents the enriched pathway/term. The size of the dots represents the number of genes, and the color indicates the –log10(FDR) value (Gray represents FDR less than 7.70E-13). The enrichment ratio was defined as the observed number divided by the expected number of genes in the annotated category in the gene list. PPI networks of (B) neurogenesis GO biological process, (D) Alzheimer’s disease KEGG pathway, and (F) abnormal synaptic transmission MPO. The nodes in the network represent genes. Key node genes are indicated with gene symbols.

### *Enc1* co-expressed genes are highly expressed in GABAergic neuron

To explore which brain cells express predominantly high levels of *Enc1* and its co-expressed genes, we analyzed the single-cell sequencing data of adult mouse brains and found that these genes were specifically expressed in three brain cell types (**Fig. 5**), Ciliated ependymal cell (205 genes), Ependymal cell_Ttr high (174 genes), and GABAergic neuron (180 genes). The GABAergic system is closely related to cognitive function, which plays an important role in neurological disorders, such as vascular dementia, AD, depression and schizophrenia[47]. Ependymal cells play a key role in cerebrospinal fluid homeostasis, brain metabolism, and removal of brain waste. These cells have been implicated in diseases throughout their life cycle, including developmental disorders, cancer, and neurodegenerative diseases[48].

**Figure 5.**
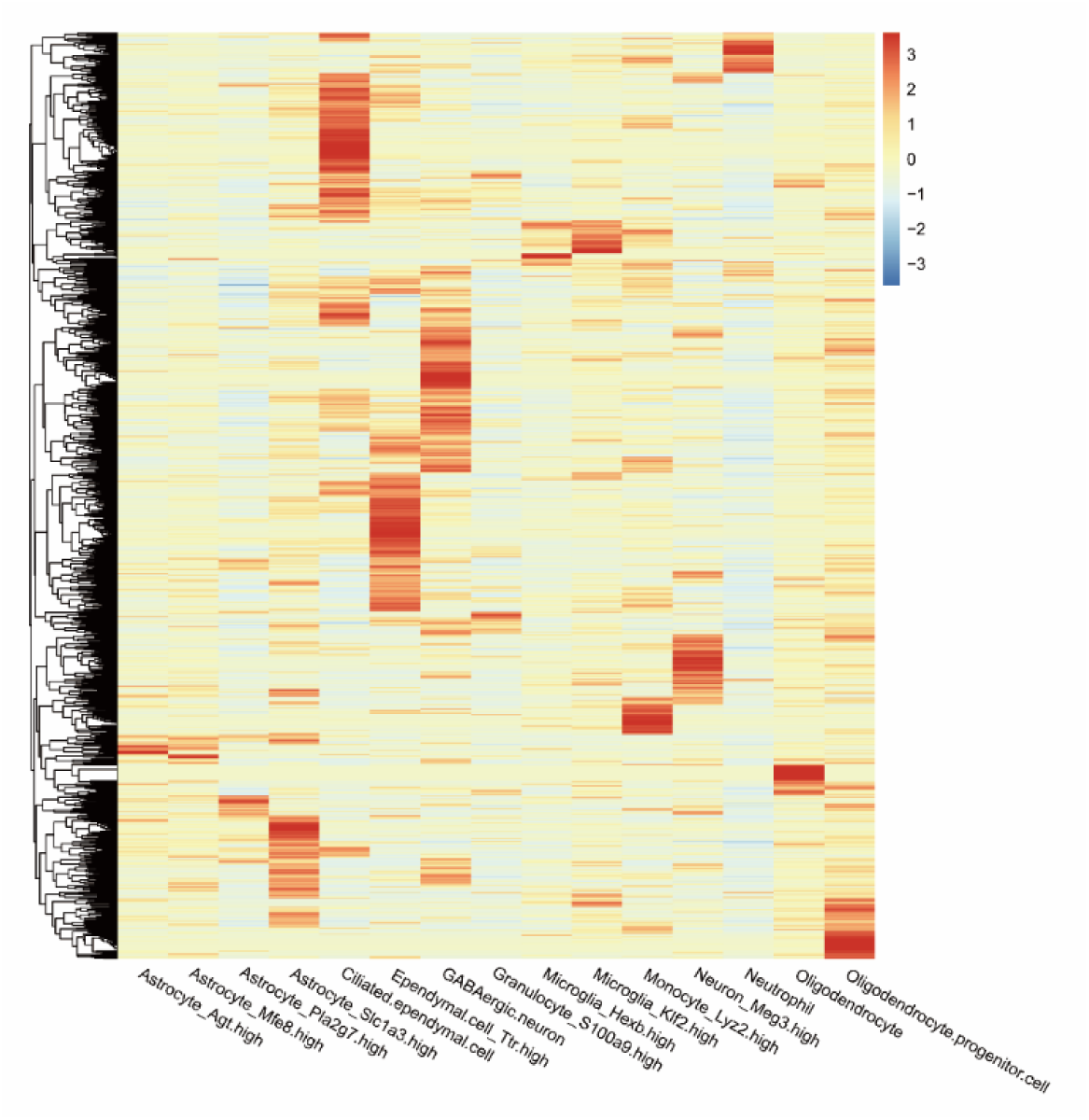
– Expression of *Enc1* and its co-expressed genes across 15 brain cell types of mice. Each column represents a cell type in the brain and each row represents a gene. The cell type specific expression data was obtained from MCA (Mouse Cell Atlas, https://bis.zju.edu.cn/MCA/).

**Figure 6.**
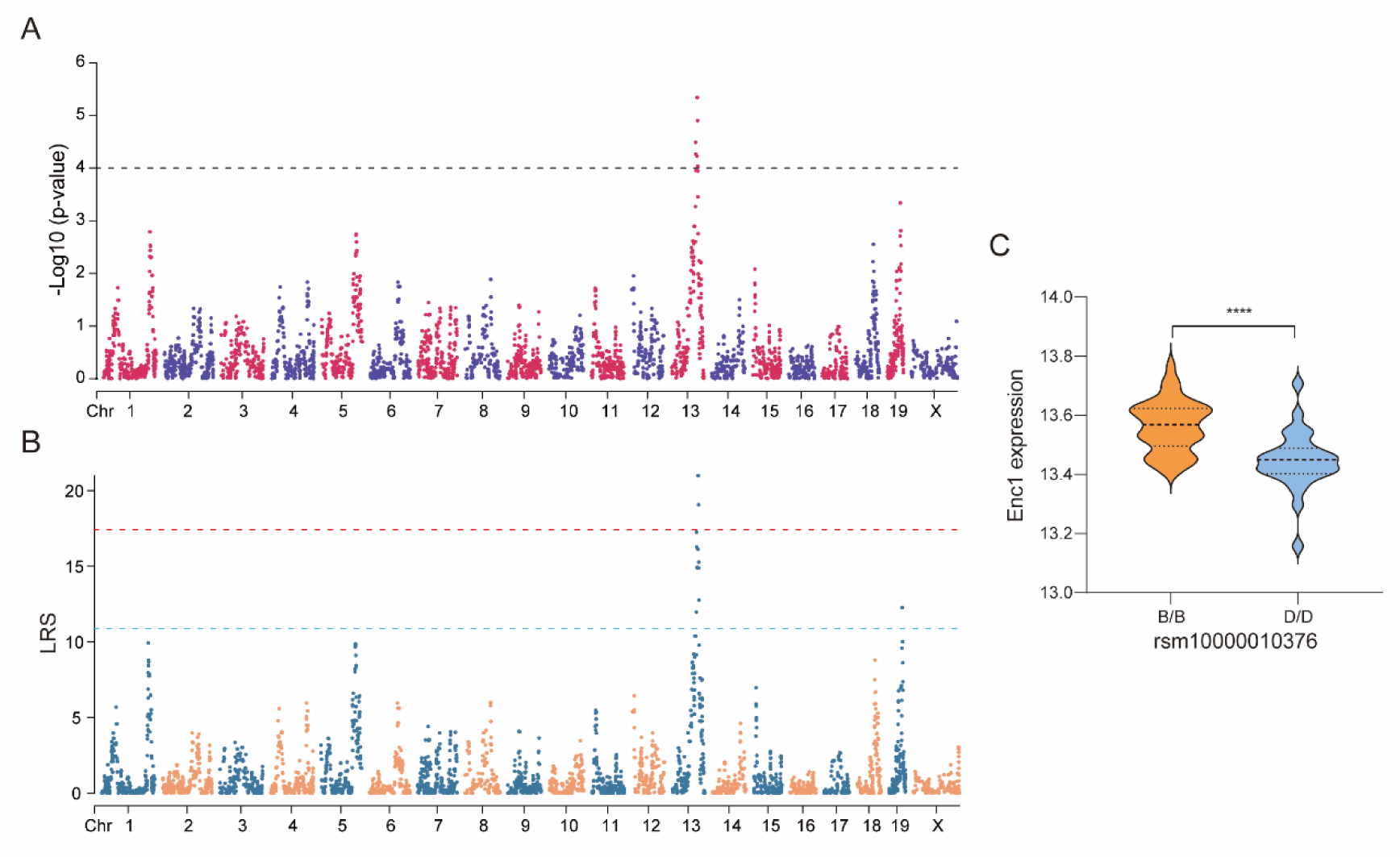
– Expression quantitative trait locus (eQTL) mapping for *Enc1*. (A). Manhattan plots showing the genome-wide regulating locus for *Enc1* identified with GEMMA, and (B) Haley-Knott Regression methods. The x-axis denotes a position on the mouse genome in megabases (Mb). The y-axis indicates the −log(p) score or likelihood ratio statistic (LRS), a measurement of the linkage between gene expression and genomic region. The grey horizontal line indicates the significant threshold of the −log(p) scores for eQTL mapping with −log(p) = 4. The red and blue horizontal lines indicated the significant and suggestive threshold of the LRS for eQTL mapping, respectively, which were 17.44 and 10.89. Genome-wide significant threshold was determined with 1000 permutation test. (C) *Enc1* was significantly different between the B and D alleles at 95.5551 Mb on Chr 13 via an unpaired t-test. ****p < 0.0001.

In addition, these genes were also significantly expressed in Astrocyte_Slc1a3 high (107 genes), Oligodendrocyte progenitor cells (94 genes), Neuron_Meg3 high (81 genes), Monocyte_Lyz2 high Neutrophil (66 genes), Microglia_Klf2 high (44 genes), and Oligodendrocyte (43 genes). Interestingly, *Enc1* gene was found to be highly expressed in microglia compared to other cell types. Molecules released from microglia have been reported to contribute to synaptic plasticity and learning and memory deficits associated with aging, AD, traumatic brain injury, and other neurological or psychiatric disorders, such as autism, depression, and post-traumatic stress disorder[49].

### The expression level of *Enc1* in the BXD hippocampus is strongly *cis*-regulated

*Enc1* is located on chromosome (Chr) 13, starting at 97.25 Mb. To investigate whether the mRNA expression of *Enc1* gene is regulated by gnomically epistatic loci, we performed genome-wide eQTL mapping, with a significance threshold of –log10(p-value) = 4. A significant eQTL for *Enc1* was mapped to Chr 13 at 95.55 Mb (rsm10000010376) with the GEMMA method (**Fig. A.6**). This was further confirmed with the Haley-Knott regression method (**Fig. B.6**). This locus was located at a distance of 0.2 Mb from *Enc1*, indicating that *Enc1* is *cis*-regulated in the BXD hippocampal region. Statistical analysis of the two cohorts (parental strains) according to the genotype of the eQTL peak position (rsm10000010376) showed that BXD mice with the B6 allele exhibited significantly higher *Enc1* expression than those carrying the D2 allele (p < 0.0001, **Fig. C.6**). *Enc1* sequence variant loci for the B6 and D6 alleles included a total of 21 missense mutations (**Table 3**).

**Table 3.** List of *Enc1* genetic variants between B6 and D2. Notes: The variant data was obtained from the MGI (Mouse Genome Informatics, https://www.informatics.jax.org/) database.

| SNP ID(GRCm39) | Position | Category | Allele<br>Summary | C57BL/6J | DBA/2J |
| --- | --- | --- | --- | --- | --- |
| rs29945732 | Chr13:97376053 | 1560 bp upstream | C/T | C | T |
| rs30008869 | Chr13:97376060 | 1553 bp upstream | C/T | C | T |
| rs29831375 | Chr13:97376255 | 1358 bp upstream | C/G | C | G |
| rs29546292 | Chr13:97376293 | 1320 bp upstream | C/T | T | C |
| rs29684293 | Chr13:97376417 | 1196 bp upstream | C/T | C | T |
| rs29546301 | Chr13:97376923 | 690 bp upstream | G/T | T | G |
| rs29566610 | Chr13:97376931 | 682 bp upstream | C/G | C | G |
| rs29246332 | Chr13:97377033 | 580 bp upstream | A/G | G | G |
| rs29534274 | Chr13:97377035 | 578 bp upstream | A/C | C | C |
| rs29590975 | Chr13:97377195 | 418 bp upstream | A/G | A | G |
| rs29238537 | Chr13:97377262 | 351 bp upstream | C/T | T | C |
| rs29920934 | Chr13:97377264 | 349 bp upstream | C/G | G | G |
| rs29620437 | Chr13:97377280 | 333 bp upstream | A/C | A | C |
| rs29818538 | Chr13:97378983 | within coordinates of | A/G | G | A |
| rs29614912 | Chr13:97379228 | within coordinates of | A/G | A | G |
| rs29516210 | Chr13:97379238 | within coordinates of | A/G | A | A |
| rs29971633 | Chr13:97382244 | within coordinates of | A/G | G | G |
| rs29225400 | Chr13:97383926 | within coordinates of | C/T | T | C |
| rs29232993 | Chr13:97383927 | within coordinates of | C/G | G | C |
| rs29551125 | Chr13:97384019 | within coordinates of | A/G | G | A |
| rs29226719 | Chr13:97384352 | within coordinates of | A/G | G | G |
| rs29714568 | Chr13:97384865 | within coordinates of | C/T | T | T |
| rs29684585 | Chr13:97385591 | within coordinates of | C/T | C | T |
| rs29529941 | Chr13:97386139 | within coordinates of | C/T | C | T |
| rs29774460 | Chr13:97387146 | within coordinates of | C/T | C | C |
| rs13465281 | Chr13:97389107 | within coordinates of | A/G | A | G |
| rs29528998 | Chr13:97389248 | within coordinates of | A/G | G | G |
| rs29551526 | Chr13:97390072 | 530 bp downstream | C/T | C | T |
| rs29571291 | Chr13:97390222 | 680 bp downstream | A/T | A | T |
| rs29730246 | Chr13:97390285 | 743 bp downstream | A/G | G | A |
| rs29735130 | Chr13:97390597 | 1055 bp downstream | A/G | G | G |
| rs29492844 | Chr13:97390979 | 1437 bp downstream | C/T | T | C |

### *ENC1* is associated with cognition in humans

*ENC1* is located on chromosome 5 (Gene ID: 8507) at 74.6 Mb in the human genome. It is highly expressed in the brain with a mean value of 179.70 ± 94.99 (**Fig. A.7**, data from NCBI, https://www.ncbi.nlm.nih.gov/). Specifically, *ENC1* is expressed in the frontal cortex, cortex and anterior cingulate cortex, and hippocampus (**Fig. B.7**, data from GTEx, https://gtexportal.org/). We also found that *ENC1* is not only cis-regulated in the human hippocampus (rs60311862, p-value = 6.00E-6), but also in brain regions of substantia nigra, amygdala, cortex, cerebellum, caudate, cerebellar hemisphere (**Fig. C.7**). In addition, GWAS revealed that the genetic polymorphisms in or nearby *ENC1* are significantly associated with cognitive-related traits (**Fig. D.7**), including age-related cognitive changes (rs76662990, p-value = 8.00E-6), PHF –tau measurement phenotype (rs7949874 / rs298387, p-value = 3E-8), and neurofibrillary tangles measurement phenotype (rs9076 / rs298387, p-value = 1.00E-12). The study shows that beta-amyloid with paired helical filaments (PHF)-tau and neurogenic fiber tangles (NFT) of P-tau were hallmarks of AD and involved in memory processes, learning and other cognitive functions with neuronal and synaptic damage[50,51].

**Figure 7.**
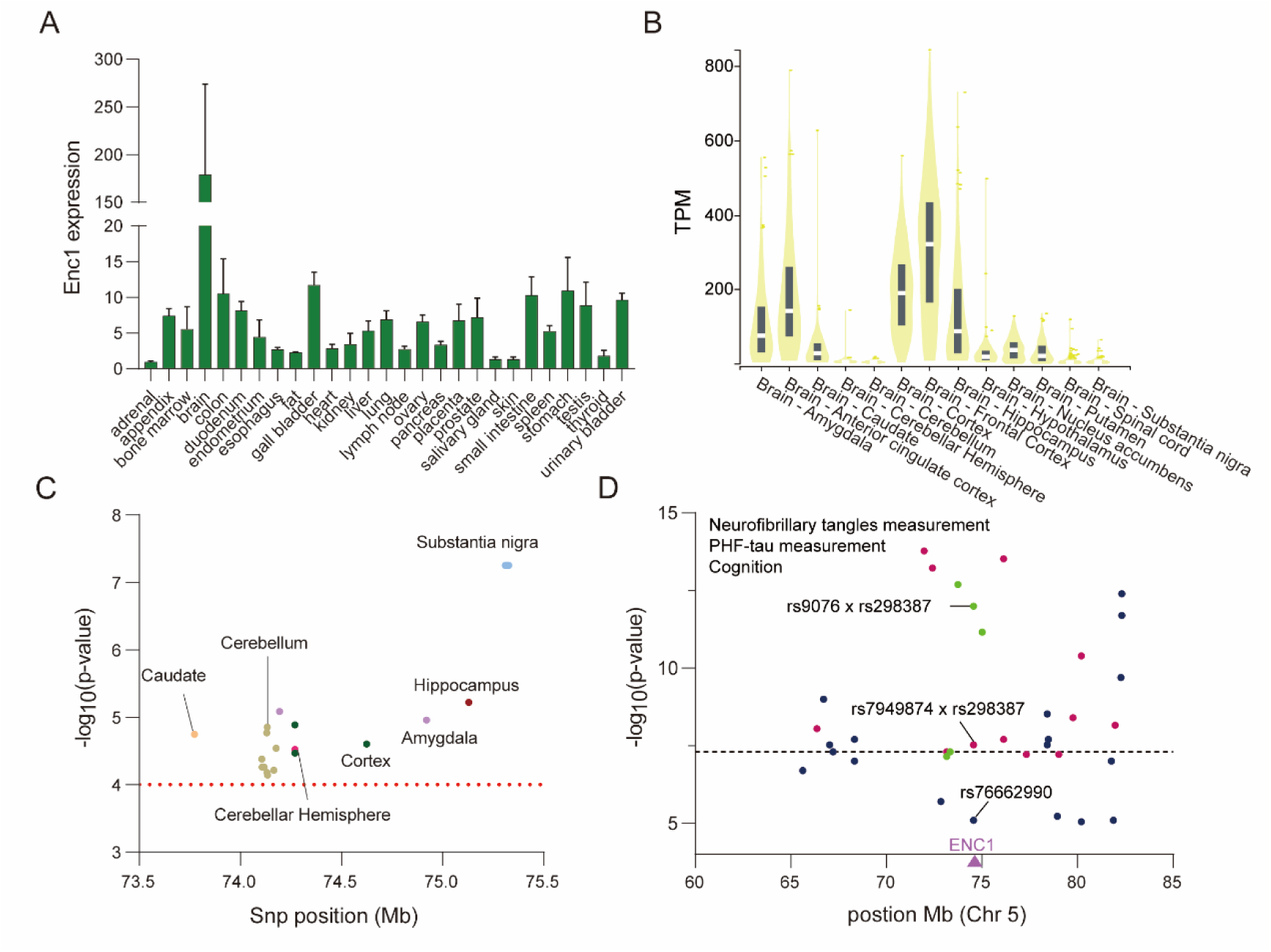
– *ENC1* gene is associated with cognition in humans. (A) *ENC1* expression in various human tissues with normalized by RPKM (Reads Per Kilobase Million). (B) *ENC1* expression in various brain regions with normalized by TPM (Transcripts Per Million). (C) Manhattan plot showing the *cis*-regulating loci of *ENC1* across the brain regions. (D) Manhattan plot showing the *ENC1* gene to be associated with cognition related traits from genome-wide association analysis. The red, blue, and green dots are associated with PHF-tau measurement, cognition, and neurofibrillary tangles measurement, respectively. The purple triangle points to the position of *ENC1*, and the threshold line is 5.00E-8. Data were obtained from the GWAS catalog.

## Discussion

In this study, we demonstrate that the hippocampal *Enc1* plays a pivotal role in regulating cognitive function. It was evidenced by the significant correlations with tens of cognitive-related traits in the BXD mice, such as y-maze and contextual fear condition performance (**Fig. B.1**), indicating impaired cognitive function with a higher level of *Enc1* expression. Supporting our results, *Enc1* knockdown significantly promotes neuronal survival and inhibits neuronal apoptosis under hypoxic/hypoglycemic injury[52]. In addition, upregulation of ENC1 adversely affects neuronal survival by downregulating cytoprotective genes, such as nuclear factor, erythroid 2-like 2 (NRF2) or by interacting with phosphorylated p62 to downregulate the autophagic pathway[12]. Healthy aging mouse models show accelerated cognitive decline in *NRF2* knockout mice relative to wild-type between 6-18 months, with cognitive deficits exacerbated at older ages (17-24 months). Furthermore, cognitive performance was noted to be enhanced after treatment with NRF2 activating compounds[53]. Our PheWAS and ePheWAS of hippocampus indicated the association of *Enc1* genetic polymorphisms with several cognitive traits in the CNS (**Fig. C-D.1**). Thus, our gene–phenotype correlation and genotype/transcriptome–phenotype association analysis demonstrated that *Enc1* is associated with cognitive function through modulation of hippocampus functions.

The *Enc1* knockdown leads to changes in levels of glucose, alkaline phosphatase, bilirubin, and cholesterol (**Table 1**). These have been reported as risk factors for cognitive impairment[54–58]. For example, high-density lipoproteins (HDL) are a heterogeneous group of lipoproteins composed of various lipids and proteins. HDL is formed in the body circulation and in the brain and may play an important role in maintaining cognitive function during aging[55]. Glucose is the main source of energy for the brain and peripheral changes in blood glucose concentration can affect cognitive performance in healthy individuals[58]. Our human GWAS results showed that *ENC1* is associated with age-related cognition, neurofibrillary tangles, and PHF-tau protein-related phenotypes in humans (**Fig. D.7**). This further suggests that Enc1 plays a role in the biological processes of cognition in both human and mouse cohorts.

Most of the co-expressed genes of *Enc1*, including the top 3 most strongly associated genes (*Arhgef12*, *Stxbp5*, *Prnp*) play a role in biological processes related to cognition. Notably, *Enc1* is positively correlated with *Prnp* (prion protein) (r = 0.626, p = 5.42E-09). The protein encoded by *Prnp* is a membrane glycosylphosphatidylinositol-anchored[59]. Mutations in this gene have been strongly associated with a variety of cognitive disorders, including Creutzfeldt-Jakob disease, Huntington’s disease-like 1, and fatal familial insomnia[60]. In addition to *Ubc*, *Ywhae* and *Mapk1*[43,44,61], our PPI network revealed several other nodal genes, including *Rnf2*, *Traf6*, *Ndn*, *Myd88*, and *Nphp1* (**Fig. 3**), which have been shown to be inextricably linked to neurological cognitive function in the brain[62–67]. *TRAF6* knockdown partially suppressed multiple inflammatory signaling pathways, reduced nerve cell apoptosis, and improved cognitive function following traumatic brain injury[65]. *Ndn* (Necdin)-deficient mice show improved spatial learning and memory in the Morris water maze[66]. In addition, *MyD88*(-/-) mice displayed impaired spatial and working memory as assessed by the Barnes maze, the water T-maze and the passive avoidance tests[67].

A gene set enrichment analysis was performed to evaluate *Enc1* pathways in the hippocampus. The results showed that the above genes mainly act in neurogenesis, synaptic signaling, CGMP-PKG signaling pathway, circadian entrainment, AD, Long-term potentiation, abnormal spatial learning and motor coordination. (**Fig. A, C, E.4**). For instance, the progressive decline in the number and maturation of newborn neurons in the hippocampus may promote or exacerbate cognitive deficits[68]. Similarly, abnormal synaptic plasticity might lead to cognitive impairments, including deficits in learning and memory, attention, and social cognition, in SCZ (Schizophrenia)[69]. CGMP-PKG signaling pathway promotes the transcription of brain-derived neurotrophic factor, which stimulates neural cell proliferation, differentiation, and survival[70]. Long-term potentiation is generally regarded as one of the main molecular mechanisms that forms the basis of learning and memory, since memory is thought to be encoded by changes in synaptic strength[71].

*Enc1* with its co-expressed genes was found to be highly expressed in GABAergic neuronal cells based on single-cell sequencing data (**Fig. 5**). Furthermore, most of the nodal genes in pathways closely related to cognition (e.g., *Mapk1*, *Bad*, *Creb1*, *Nedd4*, and *Pfn2*) were found to be highly expressed in those cells (**Fig. B, D, F.4**). GABA interneurons are major inhibitory neurons in the CNS, and they play critical roles in a variety of pathophysiological processes, including regulation of neural circuits and activity in the cortex and hippocampus, cognitive function-related neural oscillations, and information integration and processing. In addition, dysfunctional of its activity can disrupt the excitatory/inhibitory (E/I) balance in the cortex, which may be the main mechanism that can lead to cognitive dysfunction[72,73]. In addition, we found that *Enc1* is highly expressed in microglia. Microglia play a role in many aspects related to cognition, including regulating neuronal survival or death, and improving synaptic connections in the healthy brain. Moreover, microglia interact with other cells to contribute to cognitive functions. For example, neuronal activity-induced glutamate uptake and GABA release from astrocytes triggers activation of GABAB receptors in microglia. This communication pathway between neurons, astrocytes, and microglia may regulate the activity of microglia in developing neuronal networks[74]. These analyses further demonstrate the role of *Enc1* on cognitive aspects.

Through a systems analysis of *Enc1* expression and genetic regulation in the hippocampus of BXD family mice, we found that it is highly expressed in the hippocampus with 1-1.53-fold change among the strains (**Fig. A.1**). By performing eQTL analysis, we confirmed that this variation in *Enc1* expression is regulated by the local genetic variants (**Fig. A, B.6)**, where BXD mice with the B6 allele showed significantly higher *Enc1* expression than those carrying the D2 allele (**Fig. C.6**). In the human cohort also ENC1 is *cis*-regulated in the hippocampus as well as in other brain regions, including substantia nigra, amygdala, cortex, cerebellum, and cerebellar hemisphere (**Fig. C.7**). In addition, we also found the same *cis*-eQTL in several other tissues such as whole blood, lungs, spleen, thyroid, demonstrating the robustness of this *cis*-regulation in humans.

In summary, by taking advantage of the BXD family mice for genomic, phenomic, and hippocampal transcriptomic data, our results indicate that *Enc1* is associated with cognitive dysfunction-related traits. *Enc1* is involved in cognitive functions mainly through neurogenesis, synaptic signaling and CGMP-PKG signaling pathways and interacts with neurofunction related genes, *Ubc, Ywhae, Mapk1, Traf6, Ndn* and *Myd88* in GABAergic neuronal cells of the hippocampus (**Fig. 3**). Our findings demonstrate the importance of the hippocampal *Enc1* in regulating cognitive dysfunction, which may serve as a novel therapeutic target for treating cognitive dysfunction.

### Data availability statement

Raw microarray data is available on GEO (https://www.ncbi. nlm.nih.gov/geo/) under the identifier GSE84767. The normalized data is available on GeneNetwork under the “BXD” group and “Hippocampus mRNA” type with the identifier “Hippocampus Consortium M430v2 (Jun06) RMA”.

### Ethics statement

Not applicable, no human subjects.

